# Therapeutic Potential of 20E in Pulmonary Arterial Hypertension: Involvement of Mas Receptor and PI3K/Akt Pathway

**DOI:** 10.1101/2025.04.03.647130

**Authors:** Tong Lu, Weiwei Jia, Yuefei Wang, Yong Ma, Fengxia Du, Hengyu Gao, Chengrun Song, Hong Li, Xiangguo Jin, Chen Liu, Haifeng Jin, Yan Lin

**Author notes:** Corresponding author: Haifeng Jin, Heilongjiang Provincial Key Laboratory of Food & Medicine Homology and Metabolic Disease Prevention, Qiqihar Medical University, 333 Bukui Street, Qiqihar 161006, China,., Yan Lin, Heilongjiang Provincial Key Laboratory of Food & Medicine Homology and Metabolic Disease Prevention, Qiqihar Medical University, 333 Bukui Street, Qiqihar 161006, China,. Tong Lu and Weiwei Jia have contributed equally to this work.

## Abstract

**BACKGROUND:** 20-Hydroxyecdysone (20E), a natural polyhydroxylated steroid found in various edible plants, exhibits diverse pharmacological effects. This study aims to investigate the preventive and therapeutic effects of 20E on pulmonary arterial hypertension (PAH) and elucidate its underlying molecular mechanisms.

**METHODS:** A monocrotaline (MCT)-induced PAH rat model was utilized to evaluate the efficacy of 20E. The Mas receptor antagonist A779 and agonist AVE0991 were used to investigate the role of Mas in PAH progression and 20E-mediated prevention. Molecular docking and pull-down assays were conducted to confirm the interaction between 20E and the Mas receptor. In vitro, the effects of 20E on Ang II-induced proliferation and migration of human pulmonary arterial smooth muscle cells (HPASMCs) were assessed. The PI3K/Akt signaling pathway was analyzed by Western blot.

**RESULTS:** 20E prevented PAH at 30 mg/kg and 90 mg/kg, while 90 mg/kg rescued pre-existing PAH. The protective effects of 20E were attenuated by A779. 20E upregulated Mas receptor expression and directly bound to it. In vitro, 20E inhibited Ang II-induced HPASMC proliferation and migration. It also downregulated p-PI3K, p-Akt, and p-mTOR while restoring P27 and P21 expression. Furthermore, knockdown of the Mas in HPASMCs abolished the effects of 20E on these processes.

**CONCLUSIONS:** 20E inhibits PASMC proliferation and migration by activating the Mas receptor and modulating the downstream PI3K/Akt signaling pathway, thereby effectively preventing and rescuing PAH. It may be a promising pharmacological candidate for PAH treatment.

## INTRODUCTION

Pulmonary arterial hypertension (PAH), classified as group 1 and the most severe form among the 5 subtypes of pulmonary hypertension (PH), is characterized by progressive pulmonary arterial remodeling, which leads to arterial occlusion, increased pulmonary vascular resistance, and elevated mean pulmonary arterial pressure (mPAP).^1^ As the disease progresses, right ventricular (RV) hypertrophy and RV failure may occur, ultimately resulting in death.^2^ A pathological hallmark of PAH is the uncontrolled proliferation of cells within the three vascular wall layers—endothelial cells, smooth muscle cells, and fibroblasts.^3^ Notably, the excessive proliferation and reduced apoptosis of pulmonary artery smooth muscle cells (PASMCs) in the medial layer play a pivotal role in increasing vascular resistance and elevating mPAP.^4,5^ Current therapies, which include drugs that enhance the nitric oxide– cyclic guanosine monophosphate pathway, endothelin pathway antagonists, and prostacyclin pathway agonists, primarily alleviate symptoms and delay disease progression; however, a definitive cure remains elusive.^6^ ^7^

The renin-angiotensin system (RAS) plays a crucial role in maintaining cardiovascular homeostasis and consists of two primary axes: the classical angiotensin-converting enzyme (ACE)/angiotensin (Ang) II/angiotensin type 1 receptor (AT_1_R) axis and the counter-regulatory ACE2/Ang-(1-7)/Mas receptor axis.^8^ Mas is a G protein-coupled receptor encoded by the oncogene *Mas1*. During the pathological progression of PAH, the ACE/Ang II/AT_1_R axis is significantly upregulated, contributing to various deleterious cardiopulmonary effects, including vasoconstriction, cellular proliferation, oxidative stress, inflammation, and myocardial hypertrophy.^9^ In contrast, the ACE2/Ang-(1-7)/Mas axis exerts protective effects, such as vasodilation, anti-inflammatory actions, anti-proliferative effects, and anti-fibrotic effects.^10^ These opposing roles have led to increasing interest in targeting the ACE2/Ang-(1-7)/Mas axis as a potential vasoprotective strategy for PAH, cardiovascular diseases, and COVID-19.^8,9^ Accumulating evidence suggests that ACE2 or Ang-(1-7) can effectively inhibit pulmonary artery and RV remodeling in PAH animal models by activating the Mas receptor.^11–15^ Despite the potential benefits of activating the ACE2/Ang-(1-7) signaling in PAH treatment, the lack of an oral delivery system for the repeated administration of ACE2/Ang-(1-7) limits its clinical application.^16^ Thus, the development of drugs capable of directly activating the Mas receptor may offer a novel and promising approach for the prevention and treatment of PAH.

20-Hydroxyecdysone (20E), also known as β-ecdysterone, is a natural polyhydroxylated steroid with low toxicity in mammals. It is widely present in various plants, including *Spinacia oleracea*, *Chenopodium quinoa*, *Rhaponticum carthamoides*, and traditional medicinal plants like *Achyranthes* species.^17,18^ Due to its diverse biological and pharmacological properties—such as promoting protein synthesis, and exhibiting hypolipidemic, anti-diabetic, anti-inflammatory, antioxidant, and hepatoprotective effects—20E-containing dietary supplements and plant-based diets have gained increasing attention.^18,19^ For instance, studies on 20E supplementation have shown significant improvements in muscle mass, strength, and endurance without notable adverse effects.^20^ Furthermore, recent research suggests that 20E may exert beneficial effects in mammals partly through Mas receptor activation.^21,22^ Currently, a pharmaceutical grade formulation of 20E, BIO101, is undergoing clinical trials to evaluate its efficacy in treating sarcopenia (ClinicalTrials #NCT03452488) and its safety and therapeutic potential in severe COVID-19 patients (ClinicalTrials #NCT04472728).^23,24^ Despite the promising biological effects of 20E, its precise roles and mechanisms in PAH remain unclarified. This study aims to investigate the preventive and therapeutic effects of 20E in a PAH rat model and to elucidate the mechanisms by which it may inhibit pulmonary arterial remodeling.

## METHODS

### Data Availability

The detailed Materials and Methods section is available in the Supplemental Material. Data supporting the findings of this study are available from the corresponding author upon reasonable request.

### Statistical Analysis

All results are expressed as mean ±standard error of the mean (SEM). Statistical analysis was performed using GraphPad Prism 10.1.2 software. One-way analysis of variance (ANOVA) followed by Bonferroni’s post hoc test was employed to determine statistical significance, with a threshold of *p* < 0.05 considered statistically significant.

## RESULTS

### Both 20E and AVE0991 prevent the progression of PAH, and the protective effects of 20E can be blocked by A779

The dose-response study revealed that, compared to the control group, monocrotaline (MCT)-induced rats exhibited a significant increase in right ventricular systolic pressure (RVSP), accompanied by pathological changes, including pulmonary arterial remodeling and RV hypertrophy. Administration of 20E at doses of 30 mg/kg and 90 mg/kg effectively mitigated PAH progression, whereas the 10 mg/kg dose showed no significant effect (Figure S1).

Given that 20E can exert beneficial effects through Mas receptor activation,^21,22^ further experiments were conducted using the Mas receptor antagonist A779 and the agonist AVE0991 as controls to investigate the role of Mas in PAH progression and its involvement in 20E-mediated PAH prevention. The results demonstrated that both 20E and AVE0991 significantly alleviated MCT-induced increases in RVSP (Figure 1A, F), pulmonary arterial remodeling (Figure 1B, G, H), RV dysfunction, and RV hypertrophy (Figure 1E, K-N). However, the protective effects of 20E were substantially diminished in the presence of A779. Immunohistochemical analysis of α-SMA and PCNA showed that both 20E and AVE0991 markedly reduced the abnormal overexpression of these markers in the medial walls of pulmonary arteries, suggesting the suppression of excessive PASMC proliferation (Figure 1C, D, I, J). Notably, the inhibitory effect of 20E on PASMC proliferation was significantly attenuated by A779. These findings indicate the pivotal role of Mas receptor activation in countering PAH progression. Moreover, 20E effectively prevents the progression of PAH, but this preventive effect can be blocked by A779.

**Figure 1.**
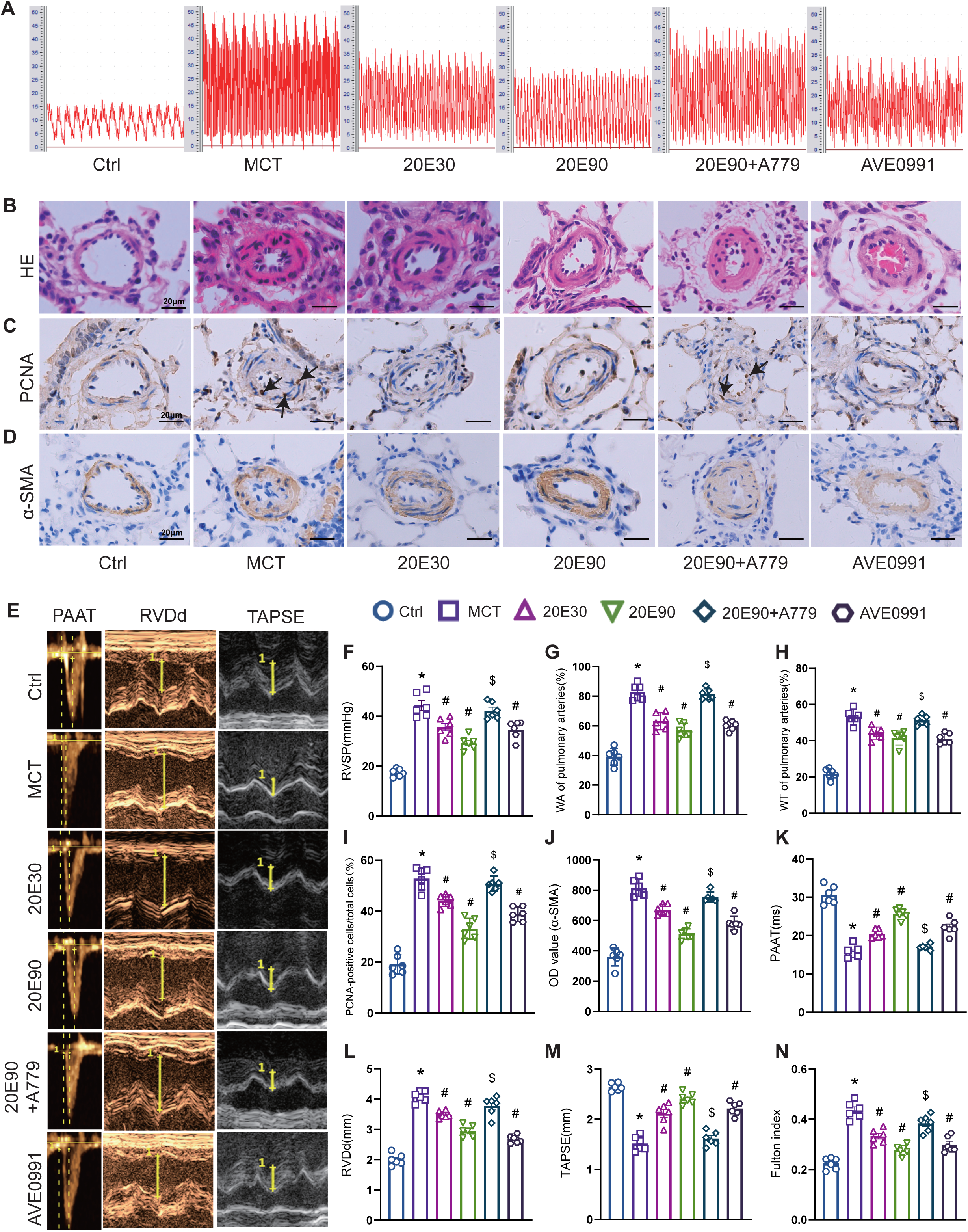
20-Hydroxyecdysone (20E) prevents monocrotaline (MCT)-induced pulmonary arterial hypertension (PAH) progression in rats. **(A)** Representative right ventricular systolic pressure (RVSP) waveforms in rats. **(B)** Hematoxylin and eosin (HE) staining of pulmonary arteries. **(C)** Immunohistochemical staining for proliferating cell nuclear antigen (PCNA) in pulmonary arteries. **(D)** Immunohistochemical staining for α-smooth muscle actin (α-SMA) in pulmonary arteries. **(E)** Representative echocardiographic images in rats. **(F)** RVSP in rats. **(G)** Medial wall area percentage (WA%) of pulmonary arteries. **(H)** Medial wall thickness percentage (WT%) of pulmonary arteries. **(I)** Percentage of PCNA-positive cells relative to the total smooth muscle cells in the medial wall of pulmonary arteries. **(J)** Quantitative analysis of the optical density (OD) value of α-SMA immunoreactivity in pulmonary arteries. **(K)** Pulmonary artery acceleration time (PAAT). **(L)** Right ventricular end-diastolic diameter (RVDd). **(M)** Tricuspid annular plane systolic excursion (TAPSE). **(N)** Right ventricle to left ventricle plus septum weight ratio (Fulton index). Data are presented as mean ± SEM (n=6). *\*P*<0.05 vs. Ctrl; *^#^P*<0.05 vs. MCT; *^$^P*<0.05 vs. 20E90.

### 20E upregulates Mas receptor expression without affecting Ang-(1-7), Ang II, or ACE2 Levels

To further investigate the effects of 20E on the ACE2/Ang-(1-7)/Mas axis, Western blotting was performed to evaluate Mas and ACE2 protein expression (Figure 2A-C), while ELISA was used to measure Ang-(1-7) and Ang II concentrations in lung tissue (Figure 2D, E). Compared to the control group, the MCT group showed a significant decrease in Ang-(1-7) levels, an increase in Ang II levels, and a marked downregulation of ACE2 protein expression, whereas Mas protein levels remained unchanged. Treatment with 20E or AVE0991 significantly increased Mas protein expression without affecting Ang-(1-7), Ang II, or ACE2 levels. Notably, the 20E-induced Mas upregulation was markedly attenuated by A779.

**Figure 2.**
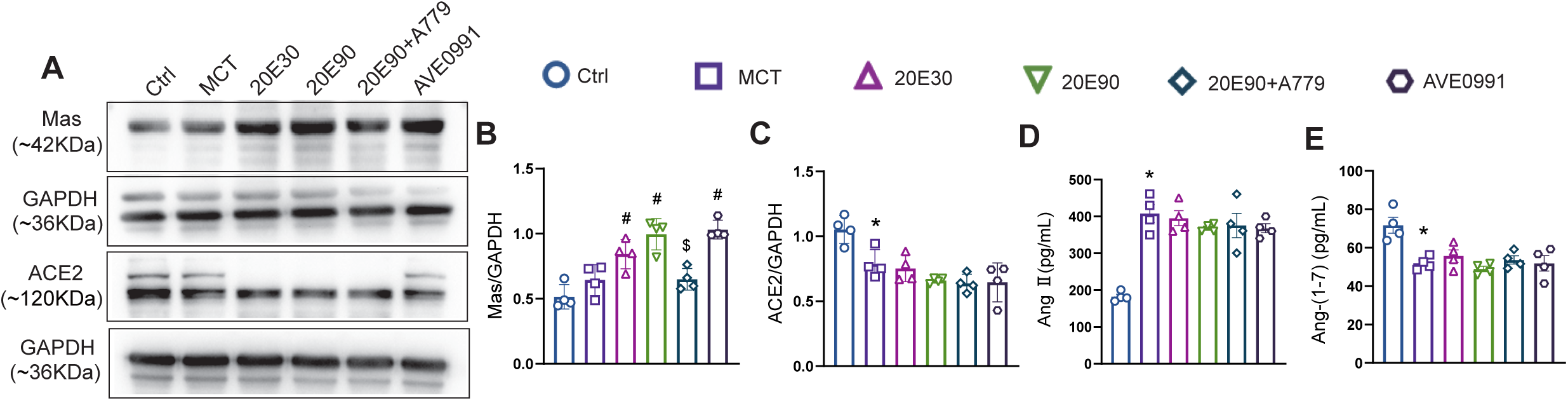
Effects of 20-Hydroxyecdysone (20E) on the ACE2/Ang-(1-7)/Mas receptor axis in rats. **(A-C)** Western blot analysis of Mas and ACE2 protein expression in lung tissues. **(D)** Ang II levels in lung tissue homogenates. **(E)** Ang-(1-7) levels in lung tissue homogenates. Data are presented as mean ± SEM (n=4). *\*P*<0.05 vs. Ctrl; *^#^P*<0.05 vs. MCT; *^$^P*<0.05 vs. 20E90.

### 20E rescues pre-existing PAH

Since PAH is not always diagnosed early, we further explored whether 20E could also rescue pre-existing PAH. Compared with the untreated MCT group, 20E at a dose of 90 mg/kg significantly reduced RVSP (Figure 3A, B), attenuated pulmonary arterial remodeling (Figure 3C, D), inhibited excessive PASMC proliferation (Figure 3F-I), improved RV function, and alleviated RV hypertrophy (Figure 3J-N). In contrast, the 30 mg/kg dose of 20E did not produce these therapeutic effects. Moreover, consistent with findings from the prevention study, 20E treatment significantly upregulated Mas protein expression in lung tissue (Figure 3O, P). Collectively, these results suggest that 20E not only prevents the development of PAH but also reverses established PAH, likely through mechanisms involving Mas receptor activation.

**Figure 3.**
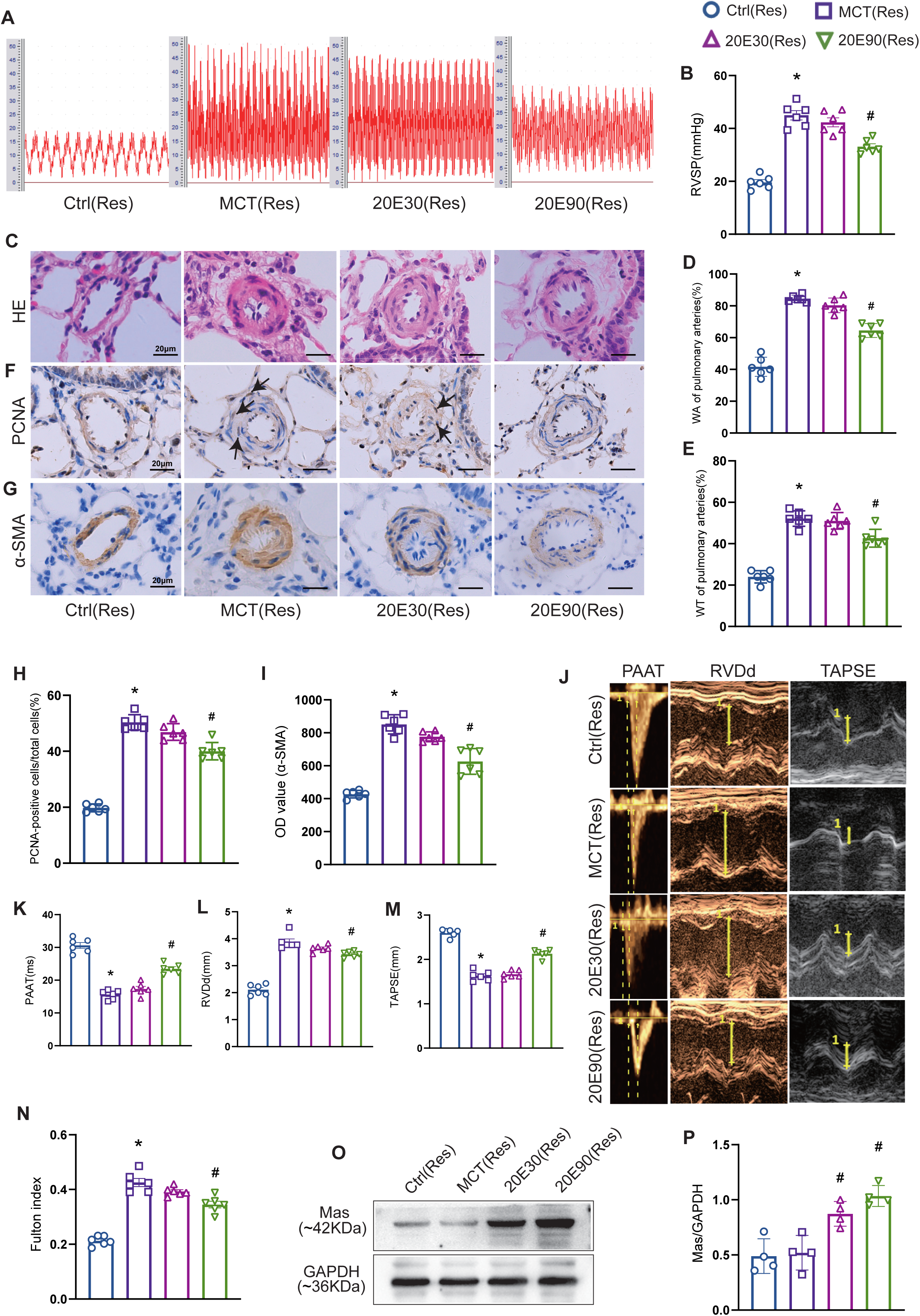
20-Hydroxyecdysone (20E) rescues pre-existing pulmonary arterial hypertension (PAH) in rats. **(A)** Representative right ventricular systolic pressure (RVSP)waveforms in rats. **(B)** RVSP in rats. **(C)** Hematoxylin and eosin (HE) staining of pulmonary arteries. **(D)** Medial wall area percentage (WA%) of pulmonary arteries. **(E)** Medial wall thickness percentage (WT%) of pulmonary arteries. **(F)** Immunohistochemical staining for proliferating cell nuclear antigen (PCNA) in pulmonary arteries. **(G)** Immunohistochemical staining for α-smooth muscle actin (α-SMA) in pulmonary arteries. **(H)** Percentage of PCNA-positive cells relative to the total smooth muscle cells in the medial wall of pulmonary arteries. **(I)** Quantitative analysis of optical density (OD) values of α-SMA immunoreactivity in pulmonary arteries. **(J)** Representative echocardiographic images in rats. **(K)** Pulmonary artery acceleration time (PAAT). **(L)** Right ventricular end-diastolic diameter (RVDd). **(M)** Tricuspid annular plane systolic excursion (TAPSE). **(N)** Right ventricle to left ventricle plus septum weight ratio (Fulton index). **(O**, **P)** Western blot analysis of Mas protein expression in lung tissues. Data are presented as mean ±SEM (A-N, n=6; O, n=4). *\*P*<0.05 vs. Ctrl (Res); *^#^P*<0.05 vs. MCT (Res).

### 20E interacts with the Mas receptor

To investigate the interaction between 20E and the Mas receptor, molecular docking analysis was performed. The results indicated a binding energy of 30 based on the London dG score and 5 according to the GBVI/WSA dG score, with a final calculated binding energy (S-value) of −7.3 kcal/mol. Visualization of the protein-ligand complex using PyMOL further confirmed this interaction, showing that the Lys146 and Arg144 residues of the Mas receptor form stable hydrogen bonds with 20E (Figure 4A). These docking results suggest that 20E exhibits a high binding affinity for the Mas receptor. To further validate the binding of 20E to Mas, we performed biotinylated protein interaction pull-down assays using human pulmonary artery smooth muscle cell (HPASMC) lysates. The results confirmed that Bio-20E specifically binds to the Mas protein in HPASMC lysates (Figure 4B, C). Collectively, these findings provide strong evidence that 20E directly interacts with the Mas receptor.

**Figure 4.**
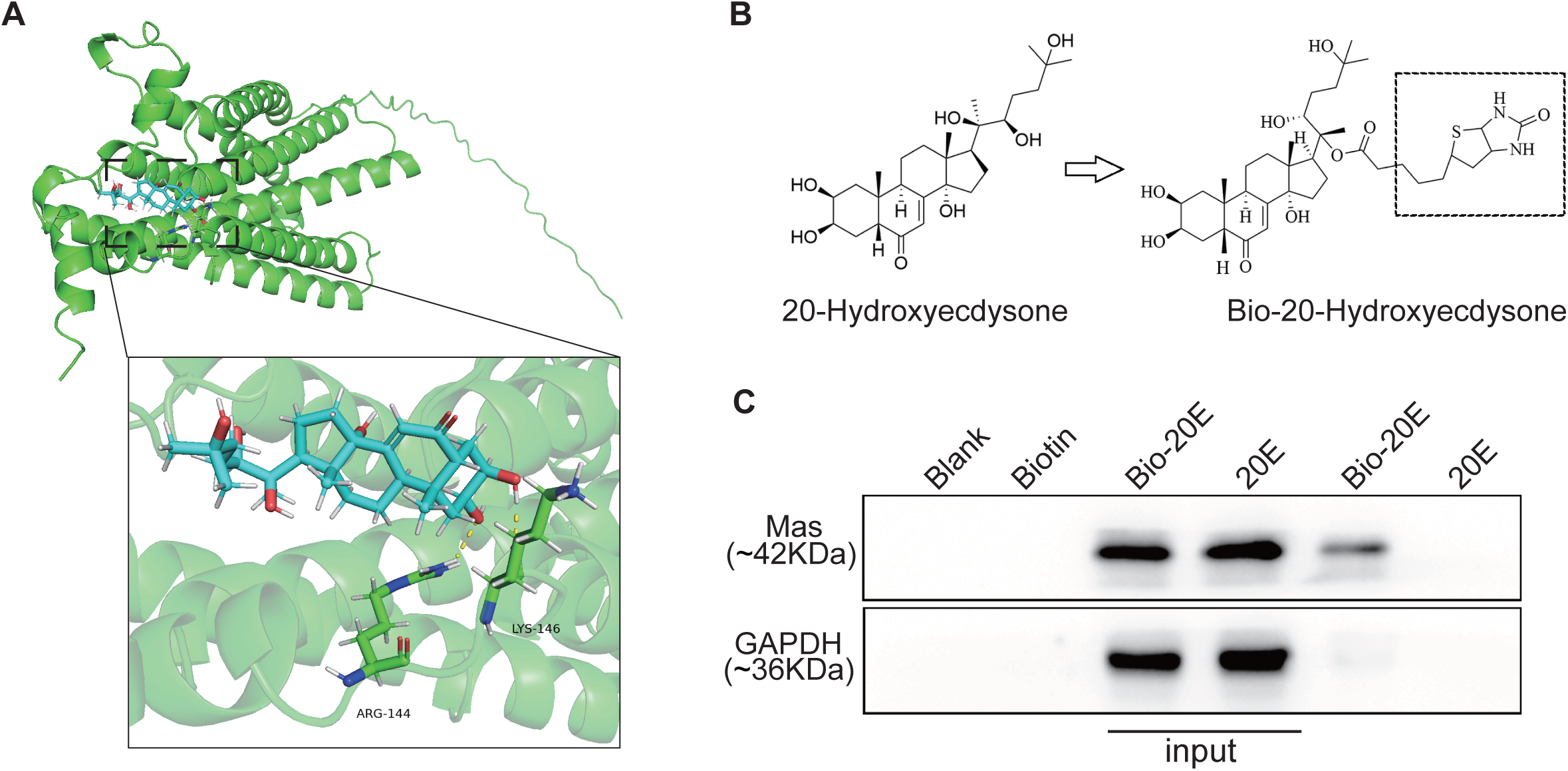
20-Hydroxyecdysone (20E) interacts with the Mas receptor. (A) Molecular docking analysis identifies two amino acid residues in Mas that interact with 20E. (B) Chemical structure of 20E and biotinylated 20E (Bio-20E). (C) Pull-down assay confirming the interaction between Bio-20E and Mas.

### 20E inhibits Ang II-induced proliferation and migration of HPASMCs and modulates the PI3K/Akt pathway

In vitro experiments were conducted to evaluate the effects of 20E on Ang II-induced abnormal proliferation and migration of HPASMCs. The results demonstrated that 20E significantly inhibited Ang II-induced HPASMC proliferation and migration in a concentration-dependent manner (Figure 5A-E). Western blot analysis further revealed that 20E markedly upregulated Mas protein expression in HPASMCs (Figure 5F).

**Figure 5.**
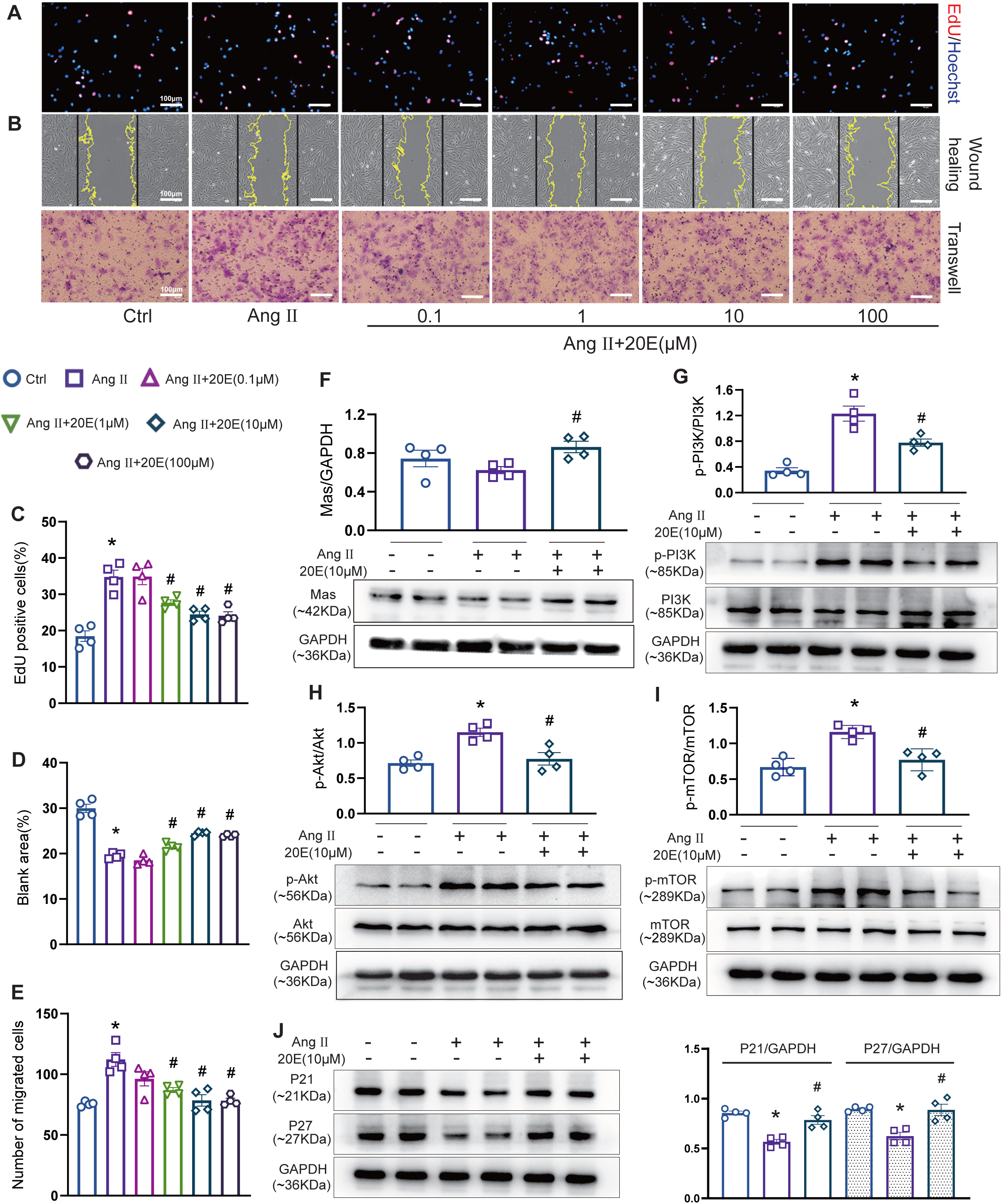
20-Hydroxyecdysone (20E) inhibits proliferation and migration of human pulmonary arterial smooth muscle cells (HPASMCs) and modulates the PI3K/Akt signaling pathway. (A) Representative EdU staining images of HPASMCs treated with different concentrations of 20E. (B) Representative images of the wound healing and Transwell migration assays in HPASMCs treated with different concentrations of 20E. (C) Quantification of EdU-positive cells. (D, E) Quantification of the wound closure area and migrated cells, respectively. (F-J) Western blot analysis of Mas, p-PI3K, p-Akt, p-mTOR, P21, and P27 protein expression in HPASMCs. Data are presented as mean ± SEM (n=4). *\*P*<0.05 vs. Ctrl; *^#^P*<0.05 vs. AngⅡ.

The PI3K/Akt signaling pathway, a crucial intracellular signaling cascade, regulates cellular processes such as survival, proliferation, and migration. Given that our previous studies identified Akt as a pivotal mediator of PASMC proliferation,^25^ we further analyzed the effects of 20E on the PI3K/Akt pathway, including its downstream target mTOR and the cell cycle regulators P27 and P21. The results showed that Ang II significantly upregulated the expression of p-PI3K, p-Akt, and p-mTOR, while markedly suppressing the expression of P27 and P21. In contrast, 20E effectively attenuated Ang II-induced upregulation of p-PI3K, p-Akt, and p-mTOR, while restoring the expression of P27 and P21 proteins (Figure 5G-J).

### Mas knockdown abolishes 20E-mediated inhibition of HPASMC proliferation and migration via PI3K/Akt pathway modulation

To further investigate whether 20E suppresses abnormal HPASMC proliferation and migration by targeting the Mas receptor and modulating the PI3K/Akt signaling pathway, we employed lentiviral transfection to knock down the Mas gene in HPASMCs (Figure 6A). Mas knockdown did not affect Ang II-induced HPASMC proliferation and migration, but it abolished the inhibitory effects of 20E on these processes (Figure 6B-F). Moreover, the regulatory effects of 20E on the expression of p-PI3K, p-Akt, p-mTOR, P27, and P21 proteins were significantly diminished in Mas-knockdown cells (Figure 6G-K). These findings suggest that 20E directly targets the Mas receptor to modulate its downstream PI3K/Akt signaling pathway, thereby inhibiting Ang II-induced HPASMC proliferation and migration.

**Figure 6.**
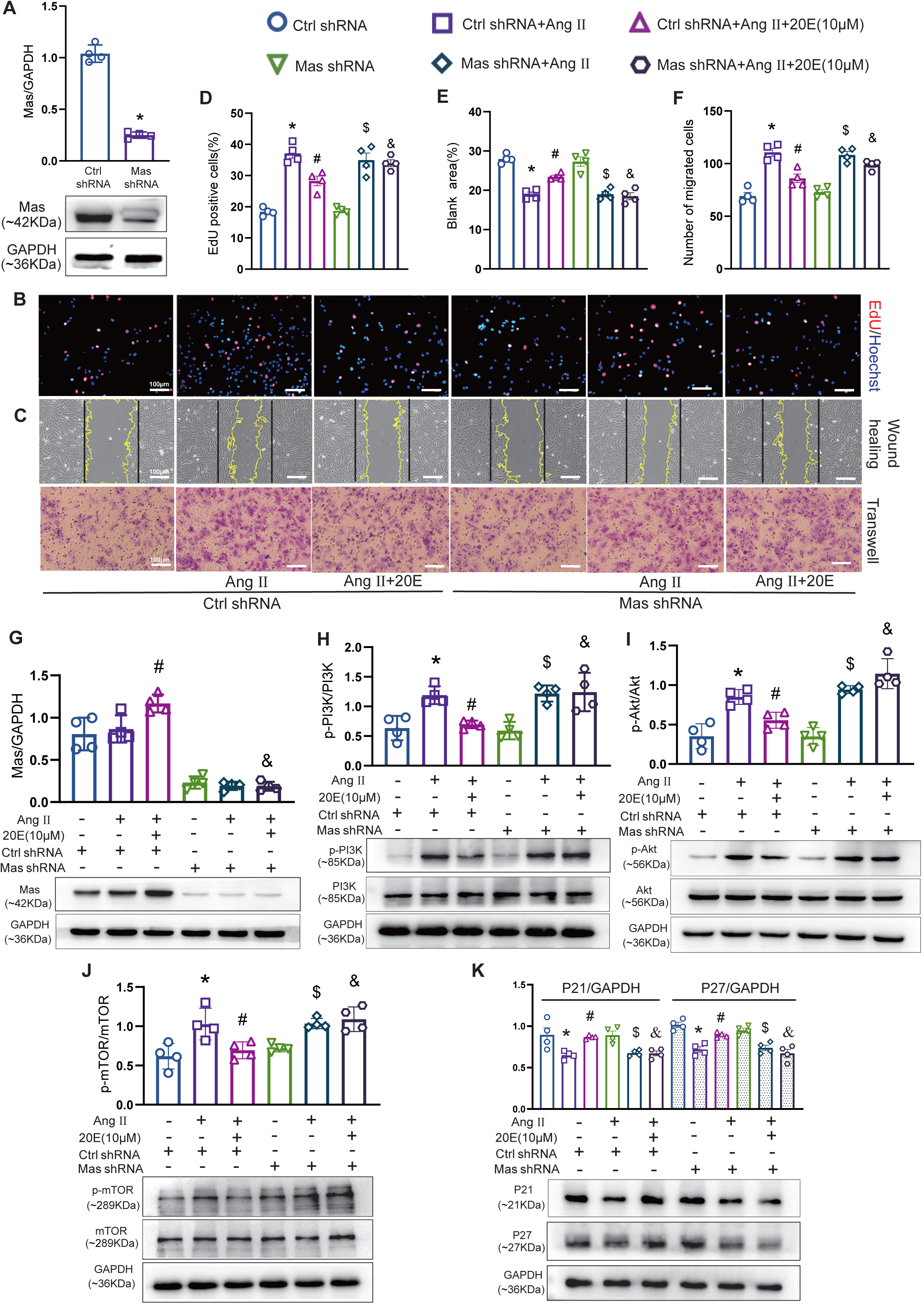
Mas knockdown abolishes the inhibitory effects of 20-Hydroxyecdysone (20E) on human pulmonary arterial smooth muscle cell (HPASMC) proliferation and migration, and impairs its regulation of the PI3K/Akt signaling pathway. (A) Western blot analysis confirming Mas knockdown in HPASMCs transfected with Mas shRNA or vector control (Ctrl shRNA). (B) Representative EdU staining images of HPASMCs following Mas knockdown. (C) Representative images of the wound healing and Transwell migration assays in Mas-knockdown HPASMCs. (D) Quantification of EdU-positive cells. (E, F) Quantification of wound closure area and migrated cells, respectively. (G–K) Western blot analysis of Mas, p-PI3K, p-Akt, p-mTOR, P21, and P27 protein expression in HPASMCs following Mas knockdown. Data are presented as mean ± SEM (n=4). *\*P*<0.05 vs. Ctrl shRNA; *^#^P*<0.05 vs. Ctrl shRNA+Ang II; *^$^P*<0.05 vs. Mas shRNA; *^&^P*<0.05 vs. Ctrl shRNA+Ang II+20E (10μM).

## Discussion

Given the complex pathogenesis of PAH and the limitations of current therapeutic options, developing more effective treatments and combination therapies has become a key research priority. The ACE2/Ang-(1-7)/Mas axis has garnered increasing attention for its potential therapeutic role in PAH.^7^ 20E, a naturally occurring compound found in various edible plants, has shown a broad spectrum of pharmacological effects in mammals. As research into its drug-like properties continues to progress, its therapeutic potential is becoming more apparent. This study demonstrates that 20E alleviates pulmonary arterial remodeling by directly binding to the Mas receptor and modulating the downstream PI3K/Akt signaling pathway.

In animal experiments, 20E exhibited significant protective effects against the onset and progression of PAH. At doses of 30 mg/kg and 90 mg/kg, 20E effectively prevented PAH development, whereas a higher dose (90 mg/kg) was required to rescue pre-existing PAH.

This stage-dependent dose requirement suggests that more advanced stages of PAH may necessitate higher doses to mitigate established pathological damage. These findings provide valuable insights into the clinical application and dose optimization of 20E.

Furthermore, similar to the Mas receptor agonist AVE0991, 20E markedly attenuated the pathological progression of PAH in rats, while significantly upregulating Mas protein expression. However, the protective effects of 20E were substantially diminished in the presence of the Mas receptor antagonist A779, highlighting the essential role of Mas receptor activation in mediating the protective effects of 20E. During the pathological progression of PAH, the protective role of ACE2 and Ang-(1-7) is primarily mediated through Mas receptor activation, which can be blocked by A779.^11,26,27^ Further analyses of ACE2, Ang-(1-7), and Ang II levels revealed that 20E did not significantly alter these factors. These findings suggest that 20E exerts its effects primarily through direct activation of the Mas receptor, rather than modulation of ACE2 or Ang-(1-7). Molecular docking studies and pull-down assays further confirmed that 20E directly interacts with the Mas receptor to inhibit the development and progression of PAH.

Due to its pathological similarities to cancer, particularly the excessive proliferation of PASMCs, PAH is recognized as a cancer-like cardiovascular disease.^28^ The abnormal proliferation of PASMCs contributes to pulmonary arterial wall thickening and luminal narrowing, which are hallmark features of pulmonary arterial remodeling and key drivers of sustained increases in mPAP and disease progression. Therefore, targeting PASMC hyperproliferation induced by pathological stimuli represents a pivotal strategy for the prevention and treatment of PAH.^29^ Immunohistochemical analyses of α-SMA and PCNA revealed that 20E significantly inhibits PASMC proliferation, an effect that was attenuated by A779. Moreover, in vitro experiments demonstrated that 20E effectively inhibits Ang II-induced hyperproliferation and migration of HPASMCs. Notably, knockdown of Mas abolished the inhibitory effects of 20E on HPASMC proliferation and migration. These findings suggest that 20E protects against pulmonary arterial remodeling by activating the Mas receptor to suppress abnormal PASMC proliferation and migration.

In the pathogenesis of PAH, the precise mechanisms by which Mas receptor activation inhibits PASMC proliferation through intracellular signaling pathways remain incompletely understood. The PI3K/Akt signaling pathway plays a pivotal role in regulating cell survival and proliferation, and is implicated in the development of both cancer and PAH.^30,31^ Activation of PI3K promotes the phosphorylation of Akt, which enhances cell survival and proliferation by activating downstream pro-proliferative targets, such as mTOR, and inhibiting cell cycle regulators, including P27 and P21.^32^ Pharmacological agents targeting this pathway, including imatinib (a receptor tyrosine kinase inhibitor) and albumin-bound rapamycin (an mTOR inhibitor), have been investigated in clinical trials for PAH treatment.^33^ This study demonstrates that 20E significantly reduces the phosphorylation levels of PI3K, Akt, and mTOR, while increasing the expression of the cell cycle inhibitors P27 and P21. However, these regulatory effects were markedly diminished in Mas-knockdown cells. Collectively, these findings suggest that 20E exerts its anti-proliferative and anti-migratory effects on PASMCs by modulating the PI3K/Akt signaling pathway through Mas receptor activation.

To the best of our knowledge, this study provides the first definitive evidence that 20E not only prevents the development of PAH but also rescues pre-existing PAH by activating the Mas receptor and inhibiting the downstream PI3K/Akt signaling pathway. However, further investigation is required to fully elucidate the mechanisms underlying the effects of 20E. First, RV failure is a major cause of mortality in PAH patients, accounting for approximately 70% of PAH-related deaths.^34^ Although this study demonstrated that 20E improves RV dysfunction and reduces RV hypertrophy, the precise mechanisms of its protective effects and its therapeutic potential in established RV failure remain unclear and require further exploration. Second, the pathogenesis of PAH is highly complex, involving multifactorial interactions, including pulmonary artery endothelial cell (PAEC) injury and dysfunction, as well as aberrant inflammatory responses. Previous studies suggest that the Mas receptor plays a pivotal role in protecting endothelial cells from injury and suppressing inflammation.^7,9,35^ Whether 20E regulates PAEC injury and inflammatory processes through the Mas receptor remains to be investigated. Addressing these questions will provide more robust evidence for the development of 20E as a promising pharmacological candidate for PAH treatment.

### PERSPECTIVES

PAH remains a challenging condition to treat due to its complex pathogenesis and the limited efficacy of current therapies. Despite advances in management, a definitive cure is still lacking. This study demonstrates that 20E, a natural compound, inhibits PASMC proliferation and migration by activating the Mas receptor and modulating the downstream PI3K/Akt signaling pathway, thereby effectively preventing and rescuing PAH. These findings suggest that 20E may serve as a promising therapeutic candidate for PAH. Future research should further investigate its effects on PAEC injury, inflammatory responses, and its therapeutic potential in RV failure.

## Sources of Funding

This work was supported by the Qiqihar Academy of Medical Sciences (2021-ZDPY-001, QMSI2022M-02), the National Natural Science Foundation of China (82074148), the Heilongjiang Provincial Natural Science Foundation (PL2024H260), the Heilongjiang Postdoctoral Science Foundation (LBH-QY22002), the Science and Technology Bureau of Qiqihar (LSFGG-2023045), and the Construction Project of Dominant Characteristic Disciplines of Qiqihar Medical University (QYZDXK-013).

## Disclosures

None

## Supplemental Material

Materials and methods

Figure S1

## Nonstandard Abbreviations and Acronyms

20E: 20-Hydroxyecdysone
PAH: pulmonary arterial hypertension
PASMC: pulmonary arterial smooth muscle cell
mPAP: mean pulmonary arterial pressure
RV: right ventricular
MCT: monocrotaline
RVSP: right ventricular systolic pressure

## NOVELTY AND RELEVANCE

### What Is New?

- 20E, a natural polyhydroxylated steroid, not only prevents the development of PAH but also rescues pre-existing PAH by activating the Mas receptor.
- 20E directly interacts with the Mas receptor, inhibiting PASMC proliferation and migration by modulating the PI3K/Akt signaling pathway.
- Knockdown of the Mas abolishes the inhibitory effects of 20E on PASMC proliferation and migration.

### What Is Relevant?

- 20E effectively reduces pulmonary artery pressure, attenuates pulmonary arterial remodeling, improves RV function, and alleviates RV hypertrophy in a PAH rat model.
- By targeting PASMC hyperproliferation, a key driver of pulmonary arterial remodeling, 20E mitigates a critical aspect of PAH pathology.
- This study identifies the Mas receptor as a key mediator in PAH progression and as a critical target for 20E’s protective effects.

### Clinical/Pathophysiological Implications

This study suggests that 20E may serve as a promising therapeutic candidate for PAH by targeting the Mas receptor and inhibiting the downstream PI3K/Akt signaling pathway. Future clinical investigations are required to confirm its efficacy and safety.

## Notes

### Competing Interest Statement

The authors have declared no competing interest.

